# Methods for detecting co-mutated pathways in cancer samples to inform treatment selection

**DOI:** 10.1101/082552

**Authors:** Tingting Jiang, Uri Shaham, Fabio Parisi, Ruth Halaban, Anton Safonov, Harriet Kluger, Sherman Weissman, Joseph Chang, Yuval Kluger

## Abstract

Tumor genomes evolve through a selection of mutations. These mutations may complement each other to promote tumorigenesis. To better understand the functional interactions of different processes in cancer, we studied mutation data of a set of tumors and identified significantly co-mutated pathways. Fisher’s exact test is a standard approach that can be used to assess the significance of the joint dysregulation of pathways pairs across a patient population. We developed a robust test to identify co-occurrence using DNA mutations, which overcomes deficiencies of the Fisher’s exact test by taking into account the large variability in overall mutation load and sequencing depth. Applying our method to a study of six common cancer types, we identify enrichment of co-mutated signal transduction pathways such as IP3 synthesis and PI3K and pairs of co-mutated pathways involving other processes such as immunity and development. We observed enrichment of clonal co-mutation of the proteasome and apoptosis pathways in colorectal cancer, which suggests potential mechanisms for immune evasion.

## Introduction

In some cancers, tumor cells sequentially acquire a set of deleterious mutations, which contribute to tumor progression(1–3). Currently, very few pairs of co-mutated genes and the order in which they are altered during tumorigenesis and tumor progression are known. In colorectal cancer, mutations in the driver genes APC, KRAS, and PIK3CA are accumulated sequentially and provide selective growth advantage over normal epithelial cells(4,5). However, it is yet unknown for most cancers whether any mutation would affect the likelihood of appearance of co-mutations in other genetically interacting genes or pathways and whether co-mutations of a pair of pathways would alter the biological behaviors in cancer such as organ specific metastasis and growth rate of cancers.

Masica and colleagues genotyped 238 known oncogenic mutations in 1000 cancers of 17 different types and used the Fisher’s exact test (FET) to study the association between mutations of pairs of driver genes(6). By applying FET, they found that oncogenes that activate the same pathways often occurred mutually exclusively. In addition, they discovered significant co-occurrence of mutations in KRAS and PIK3CA, which were known to interact at the pathway level. Similarly, Kandoth *et al.* collected over 3000 samples with whole exome-sequencing (WES) across 12 major cancer types, and used the FET to identify pairs of mutated genes (SMG) with significant co-occurrence or mutual exclusivity(7). They found that co-occurring or mutually exclusive mutations are usually associated with specific cancers types or tissues of origin.

There are some limitations to the aforementioned studies. First, the power to detect cooccurring mutations is limited by the sample size and inter-tumor heterogeneity(8–10). Mutations in genes of the same biological process typically confer similar effects in tumor progression(8,11,12). Therefore, we propose to aggregate mutations at the pathway level to investigate the prevalence of co-mutation of pairs of pathways. Second, the null hypothesis for FET is that two variables are independent of each other. However, in cancer genomics data, multiple sources of variation such as overall mutation load, gene length, or sequencing depth, could bias this assumption.

Here we propose a new method for detecting significant co-mutated pairs of pathways or gene-sets in cancer mutation datasets. Previously, a patient-specific method was proposed to discover significant single mutated pathways/gene-sets(13). Due to the large variability in the number of mutations between cancer samples, the importance of a mutated pathway in a given tumor is quantified by a score which takes into consideration the overall mutation load of the sample. An enriched mutated cancer pathway was detected if the aggregative score across all the patients was significantly higher than expected. Here we used the Poisson binomial distribution as a null model to detect the co-mutations of two pathways using a patient-based score for each pair of pathways. We simulated two datasets with different background distributions of overall mutation load modulated by spike in of mutations in several single pathways and co-mutated pathways and compared the performance of Fisher’s Exact test with our method. Our method was also applied to six different types of cancer from the TCGA project and we identified unique co-mutated pathways in these cancers. In colorectal cancer, we detected co-occurrence of mutations in the proteasome pathway and the extrinsic apoptosis pathway, suggesting a potential synergistic mechanism for immune evasion due to defects in the apoptosis pathway(14) and decreased neoantigen presentation resulting from defects in the proteasome pathway(15).

## Materials and Methods

### Simulated dataset

We simulated 100 datasets with 300 samples and 180 pathways under two different scenarios (Figure S1). In the first scenario, a desired number of mutated pathways in each sample was generated from a normal distribution with a mean of 30 and standard deviation of 10. In the second scenario, the number of mutations *N* was generated from a function *N=2^g^* where *g* followed a normal distribution with a mean of 4 and standard deviation of 1: *g~N*(4,1). For each scenario, we spiked in 25 pairs of co-occurring pathway mutations: the mutation frequencies *P(A)* of the first pathway in each pair were 0.1, 0.15, 0.2, 0.25 and 0.3 and the conditional mutation frequencies of the second pathway in each pair given the first pathway, *P(B|A),* were 0.1, 0.3, 0.5, 0.7 and 0.9, respectively. In addition to mutations in the above 25 pairs of co-mutated pathways, 14 other pathways were singly mutated with frequencies of 0.05, 0.05, 0.1, 0.1, 0.13, 0.13, 0.16, 0.16, 0.2, 0.2, 0.25, 0.25, 0.3, and 0.3, respectively. Finally, random numbers of mutations were assigned with equal probability to 180 pathways to fill out the total desired number of mutations in each sample. For each sample for which the number of single and co-mutated pathways exceeded the total desired number, we randomly turned mutated pathways into wildtype to reduce the numbers of mutated pathways until the desired numbers were reached.

### TCGA dataset

We downloaded the somatic mutations calls of 1284 tumor samples from six types of cancer including uterine corpus endometrial carcinoma (UCEC N=224), colon/rectum adenocarcinoma (COAD/READ n=224), stomach adenocarcinoma (STAD n=151), skin cutaneous melanoma (SKCM n=253), lung adenocarcinoma (LUAD n=230), and lung squamous cell carcinoma (LUSC n=178) from the Synapse workspace syn1729383 (<u>https://www.synapse.org/#!Synapse:syn1729383</u>)(7). Each somatic mutation was annotated by a C score developed by Combined Annotation Dependent Depletion (CADD)(16) (<u>http://cadd.gs.washington.edu/score</u>).

We downloaded transcriptional profiles from the TCGA data portal (<u>https://tcga-data.nci.nih.gov/tcga/</u>). We studied expression levels in terms of transcripts per million (TPM) generated using the RSEM algorithm from the RNAseqV2 data archive files. The cytolytic activity and number of predicted neoantigens of the colorectal cancer set were obtained from previous data analysis on the same TCGA samples(14).

### Pathway databases

For the analysis of the pan-cancer and colorectal cancer, we used EnrichmentMap pathway repository that consists of 2921 pathways (<u>http://download.baderlab.org/EM_Genesets/January_28_2015/Human/symbol/Human_AllPathways_January_28_2015_symbol.gmt</u>). For the development of the permutation test, we employed a subset of this repository from the BIOCARTA database. The BIOCARTA database consist 217 pathways.

### Evaluating the significance of co-mutated pathways using Fisher’s exact test

Filtered mutations were aggregated at the pathway level. We hypothesized that a pathway is dysregulated if there is at least one deleterious mutation in any of its gene members. For any pair of pathways, we used the one-sided Fisher’s exact test to determine the significance of their co-mutation. For pairs of pathways with no overlapping genes, we used the mutations in all genes of both pathways to test for co-mutations. For any pair of pathways that share common genes, we first removed the common genes and the samples mutated in the common genes, and assessed the significance of co-mutation of this pair of pathways by employing only their lists of non-overlapping genes.

### Pathway co-mutation analysis using the permutation test

To test the significance of co-mutation between any two pathways, we organized the original Yale melanoma mutation dataset in a matrix form(17,18). We simulated 1000 matrices by permuting the entries of the original data. For each matrix, we randomly shuffled the original data 100,000 times such that the number of mutations per sample and per gene remained the same as in the original data. The p-value of co-mutation of a given pair of pathways was evaluated by the fraction of these 1000 permuted matrices for which the number of co-mutations exceed the number of co-mutations in the actual (unpermuted) data.

### Co-occurrence mutation analysis at the pathway level by a patient-specific method

In this section, we describe the statistical method for assessing the significance of co-mutated pathways used throughout the other parts of the manuscript.

Let the sets G={g_i_; i=1,…,n}, S={s_j_; j=1,…,m} and Q={q_k_; k=1,…,r} represent the lists of genes, samples(patients) and pathways respectively. Let *C*_*i,j*_ be a binary random variable, getting the value *1* if gene *i* is mutated in patient *j*. We refer to our data matrix *C* as the matrix of observed values of the *C*_*i,j*_ variables, that is, *C* = (*c*_*i,j*_).

In addition, we use the indicator variable *Z*_*k,j*_, such that *Z*_*k,j*_ = 1 if sample *s*_*j*_ has a mutated gene in pathway *q*_*k*_ and *Z*_*k,j*_ = 0 otherwise. We assume that the presence/absence of mutations in pathway *q*_*k*_ in a given sample *s*_*j*_ is independent on presence/absence of mutations in this pathway in any other sample *s*_*j’*_, that is, for every pathway *q*_*k*_, *Z*_*k,j*_ is independent of *Z*_*k,j’*_, where we used *j* and *j’* to indicate indices of two different patients. Note that *Z*_*k,j*_ is a Bernoulli random variable. Denote its parameter by *p*_*k,j*_. We describe two possible approaches to estimate *p*_*k,j*_: The success probability of this Bernoulli distribution represents the probability that pathway *q*_*k*_ is mutated in sample *s*_*j*_ and denote this probability by *p*_*k,j*_. We describe two possible approaches to estimate *p*_*k,j*_:

- We first assume that probability of a single random mutation in pathway *q*_*k*_ is identical in all samples and denote this probability by φ_*k*_. The mutation probability of pathway *q*_*k*_ in sample *s*_*j*_ depends on the total number of mutations in this sample. Let *n*_*j*_ be the number of mutations in sample *s*_*j*_. Then *p*_*k,j*_ can be modeled by

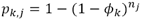 To estimate φ_*k*_, we divided the total number of mutations in pathway *q*_*k*_ by the total number of mutations in all genes across all samples.
- Estimate *p*_*k,j*_ using logistic regression. Specifically, we consider a standard logistic regression model without interaction where *p*_*k,j*_ depends on the patient *s*_*j*_ and pathway *q*_*k*_ through the coefficients α_*j*_ and δ_*k*_ respectively, *i.e.*,

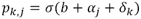

where *b* is the bias term and σ*(u)* is the sigmoid function: σ(*u*)=(1+e^−u^)^−1^.

In our experiments, we found that both approaches often perform very similarly. The results reported in this manuscript were obtained using the first approach.

Importantly, since we analyze co-mutation of pathways, for every pair *q*_*k*_, *q*_*k’*_ of pathways that share genes, the genes in the intersection must be excluded from the analysis. Hence, we modified our estimate 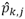 by multiplying it by the proportion of patients that had mutations outside the intersection in that pathway.

Our null model is that for every given sample *j, Z*_*k,j*_, *Z*_*k’,j*_ are independent. This implies that *Z*_*k,k’,j*_:= *Z*_*k,j*_*Z*_*k’,j*_ is also a Bernoulli random variable with parameter *p*_*k,k’,j*_:= *p*_*k,j*_*p*_*k’,j*_, being a product of independent Bernoulli random variables. Our test statistic is therefore

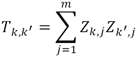

that is, the number of samples in which both pathways were mutated. Under our null model, *T*_*k,k’*_ is a sum of independent (but not identically distributed) Bernoulli random variables, also known as a Poisson-binomial random variable. Given the parameter of each Bernoulli random variable in the sum, the Cumulative Distribution Function (CDF) *F*_*k,k’*_ of a Poisson-binomial random variable can be computed iteratively(19). Let *t*_*k,k’*_ be the observed value of *T*_*k,k’*_. The p-value for the test is then obtained by

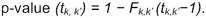

The statistical test was performed in the statistical language R using the poibin package.

### Clonal evolution analysis of co-mutated pathways

We downloaded the alignment file of Exome sequencing from cghub (<u>https://cghub.ucsc.edu/</u>) using the gtdownload. For each mutation in significantly co-mutated pathways, we extracted the reads covering the genomic regions associated with the mutations and quantified the allele frequency using samtools 2.1.13. The R package Sciclone(20) was used to infer clonality by clustering variants of similar mutant allele frequency in a single sample. Here we only selected mutations with minimum depths of 10 and mutant allele frequency of less than 0.6 to exclude mutations in CNV regions. Variants were clustered by a Bayesian binomial mixture model in Sciclone and each cluster represents one separate clone in the tumor.

### Statistical analysis

Pearson correlation coefficient was calculated between p-values generated by the Fisher’s exact test with the empirical p-values computed using the permutation test. The correlation between the patient-specific method and the empirical p-values was computed in the same way. Wilcoxon rank tests were used to compare characteristics (CYT, number of neoantigens, FASLG expression) between samples of four different conditions, namely, mutations in both pathways, only in one pathway, only in the other pathway, or in neither of the pathways.

## Results

### Workflow of patient-specific analysis to identify significantly co-mutated pathways

We developed a patient-specific method to detect enrichment of co-mutated pathways. For each patient, we focused on deleterious mutations in the coding regions. We assigned each mutation a deleteriousness score (C-score) generated by the Combined Annotation–Dependent Depletion (CADD) method(16). Single nucleotide mutations with a C-score ≥ 20 and all indels were incorporated in the enrichment analysis (Figure 1A). Then we aggregated the deleterious mutations in each patient for 2921 pathways collected from the EnrichmentMap (<u>http://baderlab.org/GeneSets</u>) (Figure 1B). To detect significantly co-mutated pathways, we investigated cancer cohorts with sample size > 200 and filtered out pathways based on the following three criteria. Pathways that were mutated in less than 20 samples (first criterion) or above 50% of the entire cohort (second criterion) were excluded from the pathway co-mutation analysis. The rational to remove pathways with few mutations across the cohort population is due to their insignificant co-mutation with other pathways. A pathway that is prevalently mutated (>50%) across the cohort population is significantly co-mutated with numerous other pathways and hence it is not indicative which of these co-mutated pathways might have synergetic interactions with this highly mutated pathway. Finally, we removed pathways whose estimated and observed mutation frequency (third criterion) have substantial deviation from each other (see Methods) such that the estimated mutation frequency was <75% of the observed mutation frequency (Figure 1C). Under the assumption that mutation co-occurrences in a given pair of pathways are independent, the number of samples for which this pair is co-mutated follows a Poisson binomial distribution. For pathways that pass the above mentioned three filtering criteria (shown in black dots in Figure 1C), we computed the significance of pathway co-mutation for any pair of pathways. This computation takes into consideration the deviation of the observed number of samples with mutations in both pathways relative to the expected number of pathway co-mutation under the null hypothesis (for details see Methods). Furthermore, for pathways that share a subset of genes we implemented an additional criterion for computing pathway co-mutation. Specifically, we excluded the overlapping genes and removed the samples that have mutations in these overlapping genes (Figure 1D). Finally, we visualized the pairs of significantly co-mutated pathways (Figure 1E) and investigated their clonal co-occurrence in each sample.

**Figure 1.**
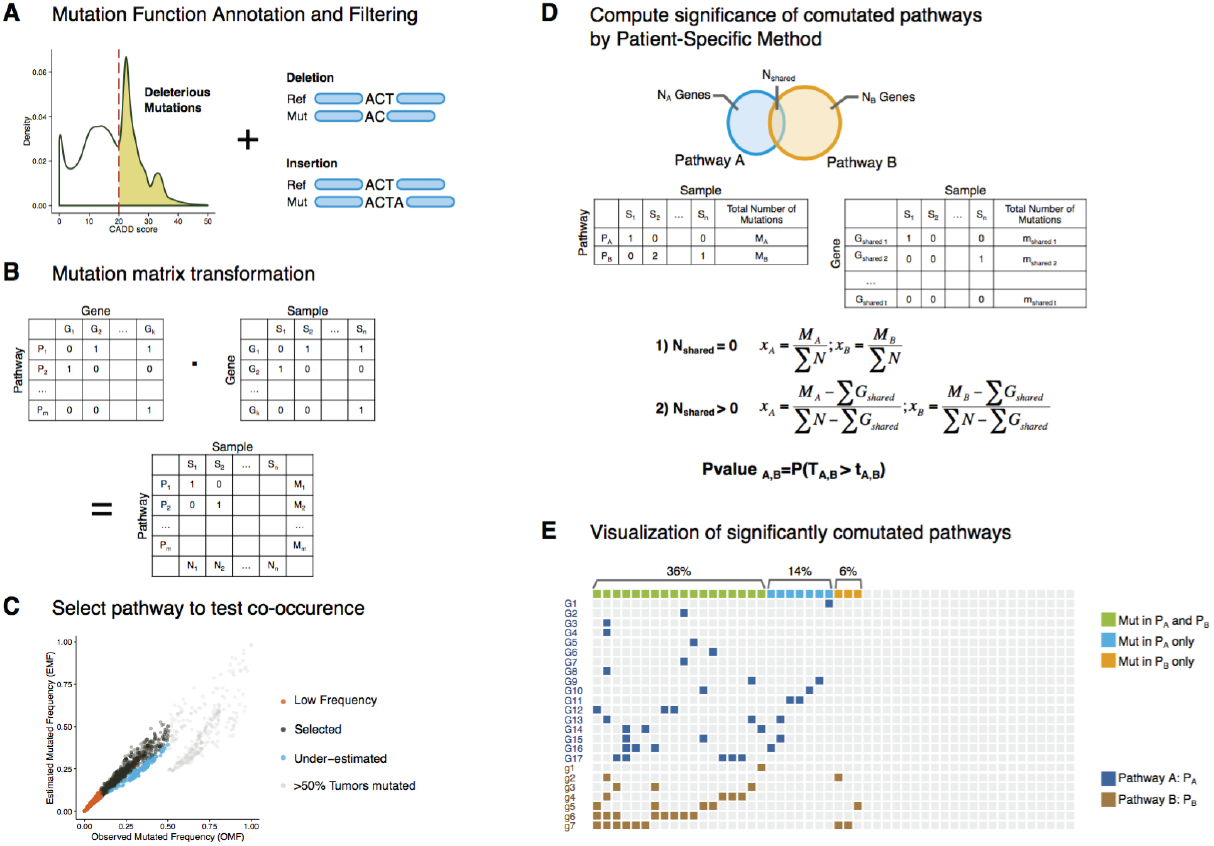
Workflow for identifying co-mutated pathways. A) Retain deleterious somatic SNV with CADD score ≥ 20 and somatic indels. B) Matrix representation for aggregating deleterious mutations into mutations at the pathway level C) Pathways represented by black dots were retained for the co-mutated pathway analysis. We filtered out pathways if they were: i) mutated in less than 20 samples (red), ii) mutated in above 50% of the samples in the cohort (grey), and iii) estimated mutation frequency was ≥25% below the observed mutation frequency (blue). D) Mutation probability of each pathway was estimated by the ratio between the mutation frequency of the pathway across the cohort and the total mutations in the cohort. For any pair of pathways (represented as pathway A and pathway B) we removed mutated genes common to both pathways (N_shared_), as well as samples with mutations in these overlapping genes. E) Visualization of significantly co-mutated pathways.

### Comparison of patient-specific method with Fisher’s exact test

Fisher’s exact test (FET) is a standard statistical test for assessing the independence of events including co-occurrence of mutations in cancer. Analyzing whole-exome sequencing data from a cohort of 303 melanoma samples from the Yale SPORE in Skin Cancer project(17,18), we observed a large dispersion in the distribution of total number of mutations across samples (with mean of 273 and standard deviation of 394, Figure 2A). The 10th and 90th percentiles of total number of mutations for this cohort are 10 and 656, respectively. We assume that a pathway is dysregulated in a given sample if at least one of its genes is deleteriously mutated and refer to this pathway as a mutated pathway. We applied the one sided Fisher’s exact test and our patient-specific statistical test to investigate the significance of co-mutations between all pairs of pathways in the BIOCARTA database, which consists of 217 pathways. The p-value distribution of our test was nearly uniform while that of FET was over-inflated with p-values smaller than 0.05 for over 93% of the pairs (Figure 2B). To further investigate which test is more representative of the actual enrichment, we generated 1000 permuted datasets to calculate the empirical p-value of co-occurring mutations in all pairs of pathways. In each permuted dataset, we started from the original mutation matrix of all the samples by all the genes and then randomly shuffled the mutations 100,000 times while preserving the same number of mutations in each gene as well as in each sample. Subsequently we determined the empirical p-value of co-mutation of any pair of pathways. For each pair of pathways, we computed the fraction of these 1000 permutated datasets in which the number of samples with co-mutations for this pair of pathways is larger compared to the number of samples with co-mutations for the observed (un-permuted) data (Figure S2). We compared the empirical p-value from the permutation test with that of the patient-specific method and FET respectively (Figure 2C, 2D). Interestingly as shown in Figure 2C and Figure 2D, the p-value of our test was highly correlated with the empirical p-value (R=0.88, Kendall’s tau coefficient t=0.68), while the correlation coefficient between the empirical p-value and FET p-value was much lower (R=0.15, t=0.33).

**Figure 2.**
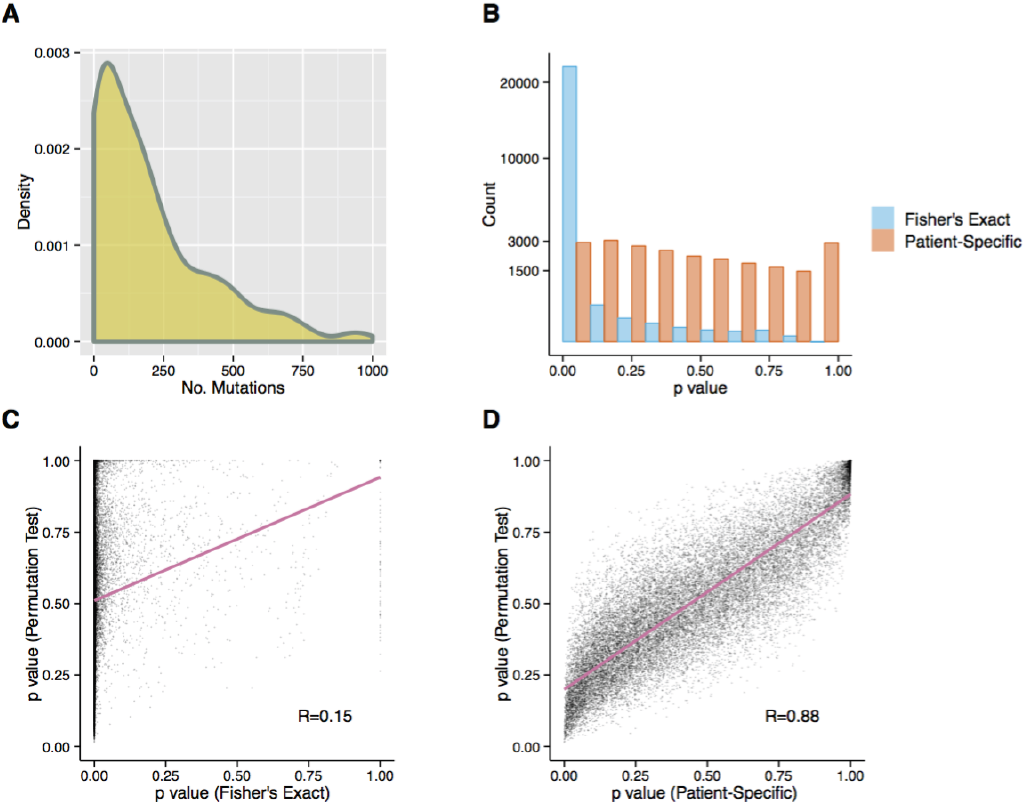
Distributions of p-values for pathway co-mutation using the patient-specific method and Fisher-exact test. A) Distribution of number of mutations in 303 Yale melanoma dataset. B) Distribution of p-values for pathway co-mutation using 217 pathways of the BIOCARTA database. C) Scatterplot of p-values of Fisher-exact test vs. p-values of a permutation test. D) Scatterplot of p-values of the patient-specific method vs. p-values of a permutation test.

### Comparison of the patient-specific test and Fisher’s exact test in simulated data

To examine the performance of the patient-specific method and the FET, we simulated two scenarios: database with two different background distributions of overall mutation load each modulated by spike in of mutations in 14 single pathways as well as 25 co-mutated pathways. The database in each of these two scenarios consisted of 100 independent simulated datasets with size of 300 samples and 180 pathways (Figure S1). In the first scenario the total number of mutations per patient (from background and spike in mutations) is normally distributed with a small dispersion (Mean=30, Sd=10, Figure 3A), and in the second scenario the total number of mutations per patient is a log-normal distribution with large dispersion (Mean=4, Sd=1, Figure 3B). Spike in mutations of the 14 singly-mutated pathways and 25 co-mutated pathways pairs were added to the background mutation as follows: For each dataset, mutations were assigned to each patient such that the singly-mutated pathways were mutated with a different mutation probability within the range of 0.05 to 0.5. Similarly, co-mutations were assigned to patients such that one pathway of each pair of co-mutated pathways (pathway A) was mutated with a different mutation probability within the range 0.1 to 0.3 and the second member of the pathway pair (pathway B) was altered with a different conditional probability (conditioned on the mutation in pathway A) within the range of 0.1 to 0.9. A detailed procedure of our simulations is provided in the Methods section. Using these simulated datasets, we compared the patient-specific method and FET in terms of sensitivity (power) and precision. The 25 spiked-in co-mutated pathways represent the positive events. Positive events with significant co-occurrence based on the statistical test of choice are referred to as true positives. Significantly co-mutated pathways in the remaining pathway pairs represent the background noise and we refer to them as false positives.

**Figure 3.**
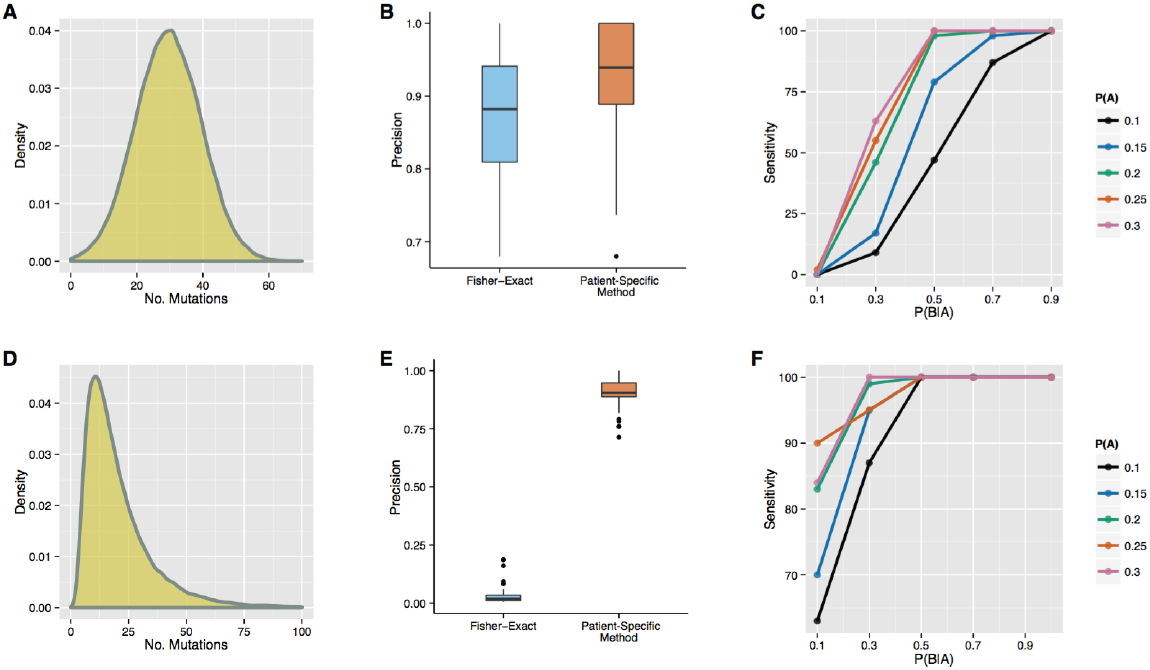
Performance of the patient-specific method and the Fisher-exact test in simulated datasets. A) Simulated mutation distribution with a small dispersion (first scenario). B) Precisions of Fisher-exact test and the patient-specific method in 100 simulated datasets (first scenario). C) Sensitivity of detecting the spiked co-mutated pathways with different mutation frequencies *P(A)* and conditional probabilities *P(B|A)* in 100 simulated datasets (first scenario). D) Simulated mutation distribution with a large dispersion (second scenario). E) Precisions of Fisher-exact test and the patient-specific method in 100 simulated datasets (second scenario). F) Sensitivity of detecting the spiked co-mutated pathways with different mutation frequencies *P(A)* and conditional probabilities *P(B|A)* in 100 simulated datasets (second scenario).

In the first scenario, the precision of the FET (median=0.88) was significantly lower than that of the patient-specific approach (median=0.93) with an fdr threshold 0.05 (paired t-test p-value=6*10^−7^, Figure 3B) over 100 simulations. In the second scenario the precision of the FET is significantly lower compared to that of the first scenario (median=0.02), while the precision values of the patient-specific method in both scenarios are similar (median=0.90). As shown in Figure 3E, the precision of the patient-specific method is significantly higher than that of the FET (paired t-test p value<2.2*10^−16^) over 100 simulations. Similar to the results shown in Figure 2B and Figure 2C, the FET does not differentiate between true and false positives in the scenario of over dispersed mutation distribution as shown in Figure 3D. Furthermore, we assessed the sensitivity of the patient-specific test to identify true co-mutated pathways with different intensity of mutation probability and conditional probability for the two scenarios with normal and over dispersed mutational load (Figure 3C, 3F). In both scenarios, the sensitivity of the patient-specific approach to detect pathway co-mutation increases as a function of the mutation probability *P(A)* and the conditional mutation probability *P(B|A)*. We observed that even for moderate values of *P(A)* and *P(B|A)* the patient-specific method reached higher values of sensitivity in the second scenario as compared with the first scenario. For example, when the conditional probability *P(B|A) = 0.5* the sensitivity to detect significantly co-mutated pathways is 100% even when the mutation probability *P(A)* is as low as *0.1*.

### Enriched co-mutated pathways in specific cancers and across multiple cancers

Mutations from TCGA were processed and filtered to obtain a data set representing 1284 tumors from 6 different cancer types including uterine corpus endometrial carcinoma (UCEC N=224), colon/rectum adenocarcinoma (COAD/READ n=224), stomach adenocarcinoma (STAD n=151), skin cutaneous melanoma (SKCM n=253), lung adenocarcinoma (LUAD n=230), and lung squamous cell carcinoma (LUSC n=178) profiled by whole-exome sequencing (Figure S3). A total of 281,979 deleterious somatic mutations in 20,817 genes were used for this analysis and 2321 out of 2921 pathways were selected using filtering steps shown in Figure 1C. The distribution of the number of mutations in each cancer type as well as in the entire cohort exhibits large dispersion (Figure S4). The number of mutations and selected pathways of each cancer type are shown in Table S1. We applied our patient-specific method to all the samples and to each cancer type to identify significantly co-mutated pathways with a p-value <5×10^−5^. Since the pathway datasets were collected from multiple sources, there was some redundancy in overlapping pathways. To eliminate these redundant pairs of pathways, we discarded from the analysis any pair for which the number of overlapping genes exceeds 50% in one or both pathways. We identified 283 significantly co-mutated pathways in a combined dataset consisting six cancers from the pan-cancer dataset^12^, and 0 - 88 significantly co-mutated pathways in these six individual cancers (Figure 4A, TableS2). Interestingly, the number of significantly co-mutated pathways in these six cohorts was not related to their sample size.

**Figure 4.**
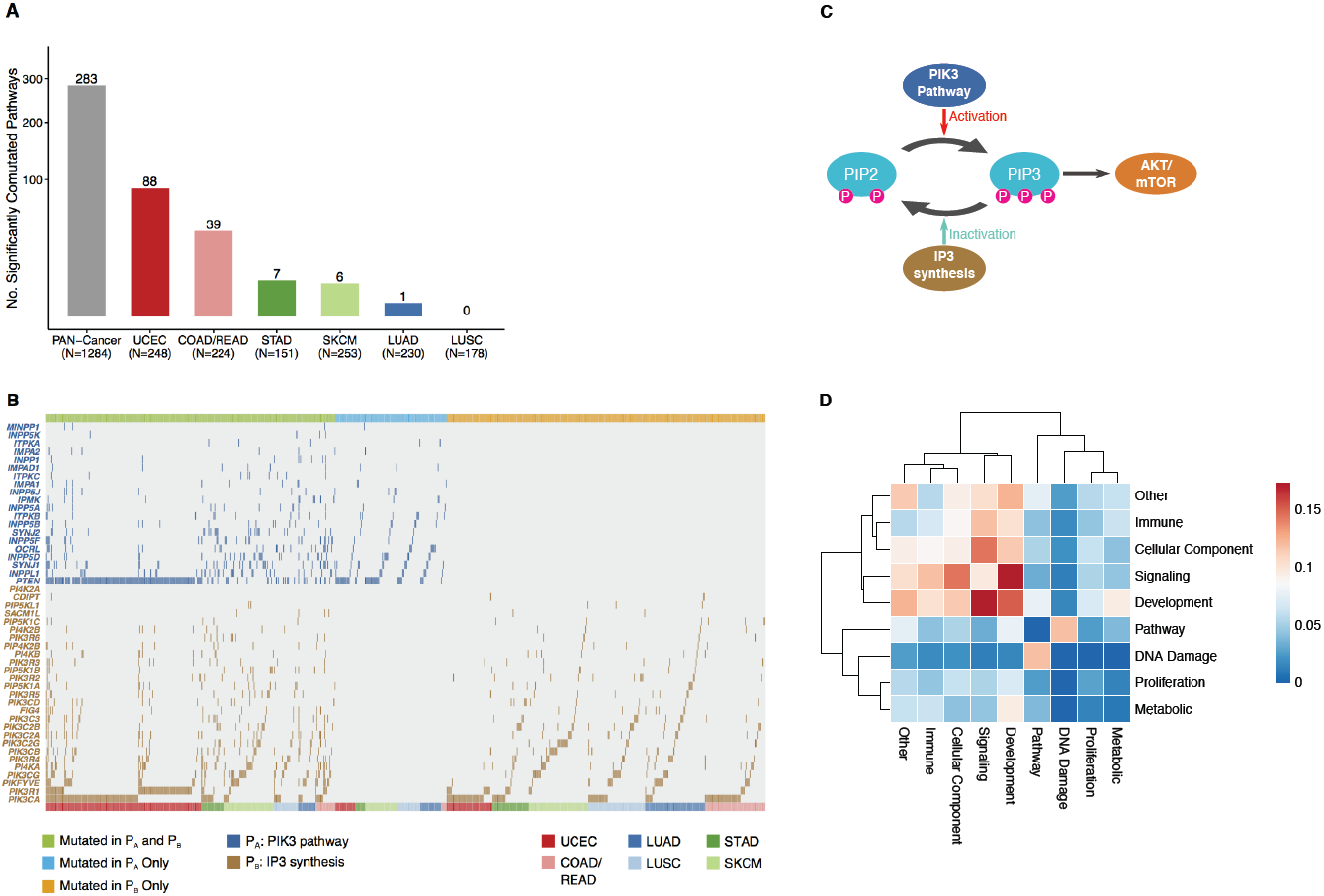
Pan-cancer and cancer type specific significant co-mutated pathways. A) Number of significantly co-mutated pathways in the pan-cancer analysis as well as in different cancer types. B) PIK3 and IP3 pathways were the most significantly co-mutated pathways in the pan-cancer analysis C) PIP3 is regulated by the PIK3 and IP3 pathways. D) Co-mutation frequency of cancer hallmarks in the pan-cancer data.

We applied the patient-specific method to the pan-cancer database and observed that the most significantly co-mutated pathways (p value=0) were IP3 metabolism (superpathway of D-myo-inositol (1,4,5)-trisphosphate metabolism) and the PI3K (3-phosphoinositide biosynthesis) pathway. Interaction between these two pathways and their relevance to cancer progression is well documented(21). Out of 1284 samples, 286 (22.3%), 110 (8.6%) and 315 (24.5%) samples had mutations in both pathways, IP3 metabolism pathway only and PIK3 pathway only respectively (Figure 4B). The most frequently mutated gene associated with the IP3 metabolism pathway was *PTEN,* which is involved in the removal of phosphate from PIP3 and hence with its degradatio*n,* and the top two most frequently mutated genes in the PI3K pathway were *PIK3CA* and *PIK3R1*. Notably, other genes in these pathways were mutated and different pairs of genes were co-mutated in different patients, indicating heterogeneity within co-perturbed pathways. The cross-talk between these two pathways involves phosphatidylinositol (3,4,5)-trisphosphate (PIP3)(22–24). Both gain-of-function mutations in the PI3K pathway and loss-of-function mutations in the IP3 metabolism pathway increases PIP3 level and therefore activates the downstream AKT/mTOR pathways, which are critical for cell survival and proliferation during cancer progression. (Figure 4C).

Another way to study co-mutated processes in cancer is by mapping significantly co-mutated pathways to eight important cancer characteristic processes termed the hallmarks of cancer(25). Pathways with ambiguous assignment to one of these eight categories were assigned to an additional category labeled “other” (Table S3). To determine the enrichment of co-mutated hallmarks of cancer, we computed the pair-wise enrichment score by the total number of significant pairs normalized by the total number of pathways in each category (Figure 4D). In the multi-tumor type analysis, pathways in immune, signaling, Cellular component and development categories were more likely to be co-mutated among themselves. The top two enrichment signals were between signaling and development categories and between signaling and cellular component categories. While co-mutations between pairs of hallmarks of cancer such as ‘DNA Damage’ and ‘Proliferation’, were less frequent.

### Co-mutation of the proteasome and extrinsic apoptosis pathways in colorectal cancer suggests an immune escape mechanism

In the TCGA colorectal cancer data set, the degradation of GLI1 by the proteasome (proteasome pathway) and the negative effector of the FAS and TNF-a pathway (apoptosis pathway) were significantly co-mutated (p-value =3.5*10^−5^). Out of 224 samples, 29 (12.9%), 24 (10.7%) and 4 (1.7%) samples had mutations in both pathways, the proteasome pathway only and the apoptosis pathway only, respectively (Figure 5A).

**Figure 5.**
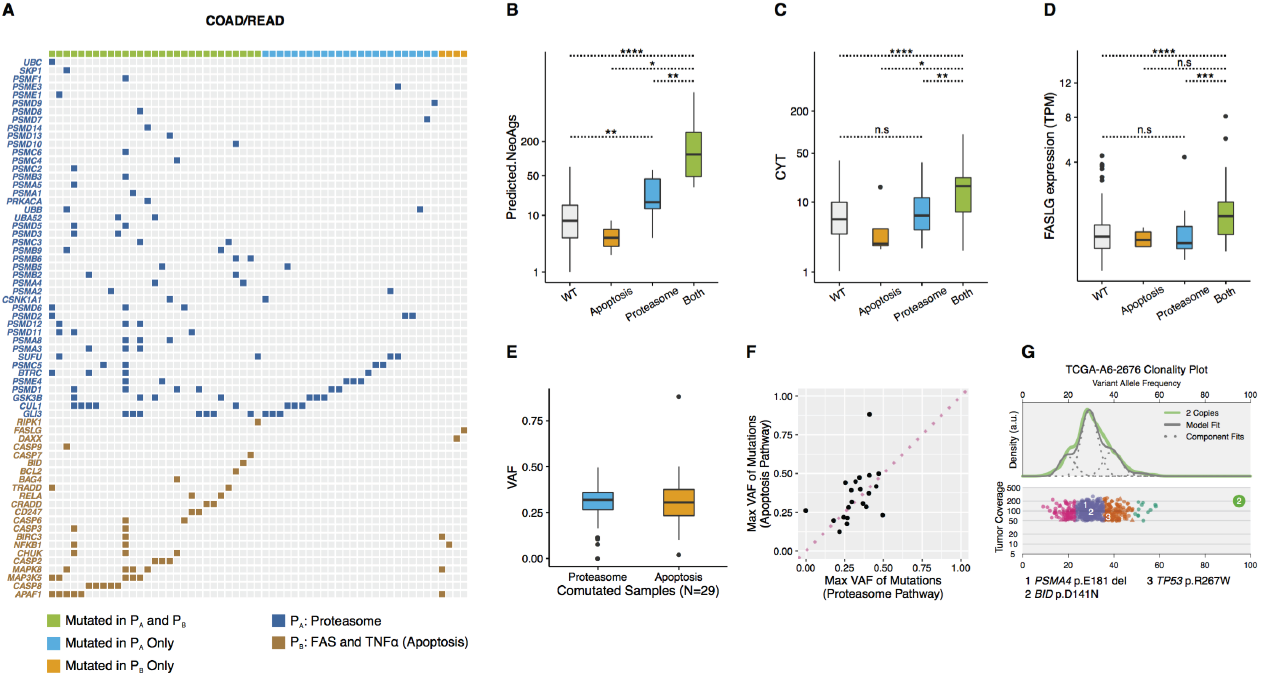
Co-mutation of the proteasome and extrinsic apoptosis pathways in colorectal cancer is associated with immune escape. A) The proteasome and the apoptosis pathways were significantly co-mutated in colorectal cancer. B) Distribution of predicted neoantigens in tumors with mutation in both pathways, in proteasome alone, in apoptosis alone, and in WT (tumor with no mutations in these two pathways). C) Distribution of cytolytic activity (CYT) in tumors with mutations in both pathways, in proteasome alone, in apoptosis alone, and in WT. D) Distribution of FASLG expression in tumors with mutation in both pathways, in proteasome alone, in apoptosis alone and in WT. E) Distribution of variant allele frequency of mutations in the proteasome and apoptosis pathways in tumors with co-mutations in these two pathways. F) Scatterplot of maximum variant allele frequency of mutations in the proteasome and the apoptosis pathways in tumors with co-mutations in these two pathways. G) Clonal analysis of sample TCGA-A6-2676. PSMA4 and BID mutations were found in the same subclone.

#### Association between mutations in the apoptosis pathway and immune escape

The apoptosis pathway includes the cancer driver gene *CASP8*, which can be mutated in cancer, resulting in immune evasion via inhibition of FasL- or TRAIL-induced apoptosis. In the proteasome pathway, different subunits of the proteasome were mutated across different samples. The patterns of mutation in both pathways were spread over different positions in different genes rather than in a small number of hotspots. This suggests that most of these mutations are loss of function mutations(7,14,26). The proteasome degrades mutated proteins and generates peptides carrying neo-antigens. We therefore hypothesized that co-mutation of the proteasome and apoptosis pathways leads to immune escape by decreasing the presentation of neoantigens and/or by decreasing the susceptibility of the cells to apoptosis.

#### Tumor sub-setting based on mutations in apoptosis or proteasome pathways and association with neo-antigens and intra-tumoral cytolytic activity

We categorized the entire cohort into four groups: (i) tumors with mutations in both pathways, (ii) tumors with mutations in the apoptosis pathway only (iii) tumors with mutations in the proteasomal pathway only and (iv) tumors without mutations in either. To explore whether these four groups are associated with distinct immune activities we applied the Wilcoxon rank test between each pair of these four groups to compare: a) the prevalence of their predicted neoantigens, b) the cytolytic activity (CYT) associated with the tumor. Tumors with mutations in the proteasome pathway alone had significantly more predicted neoantigens than the WT tumors (p-value= 0.002). However, there was no significant difference in their associated cytolytic activity (p-value=0.53), indicating that fraction of the predicted neoantigens were not presented, which could be attributed to impaired proteasome function (Figure 5B, 5C). Tumors with mutations in both pathways had significantly more predicted neoantigens (median = 118) compared to tumors with mutations in the apoptosis pathway only (median =5, p-value=0.01), in the proteasome pathway only (median=17, p-value=7*10^−4^) or with no mutations in either of these two pathways (median=8, p-value=1.2*10-^9^). Interestingly, tumors with mutations in both pathways showed higher cytolytic activity (median=16.9) than the other three groups of tumors (Figure 5B, 5C). Specifically, cytolytic activity was significantly lower in tumors with mutations in the proteasome pathway only (median=2.5, p-value=0.02), the apoptosis pathway only (median=6.3, p-value=9*10^−4^), and neither pathway (median=5.4, p-value=1.1*10^−6^). Samples with mutations in the proteosome pathway but not in the apoptosis pathway had more predicted neoantigens compared with samples that have no mutations in either pathway (WT). However, the cytolytic activity in these samples is similar. This could be attributed to the mutations in proteasome genes that prevent the proteasome to properly present the neoantigenes.

Seeing that cancer cells can potentially evade FasL induced apoptosis by acquiring mutations in the apoptosis pathway, we studied the association with mutations in the proteasome and apoptosis pathways. Tumors with mutations in both pathways exhibited significantly higher expression of FASLG compared to proteasome only (p-value=5*10^−4^) and WT group (p-value=9*10^−6^ Figure 5D). There was no difference in FASLG expression between the tumors in the group that has no mutations in this pair of pathways and the group that has mutations only in the proteasome pathway (p-value=0.62). We therefore hypothesize that tumors with mutations in both pathways can evade the immune system by either reducing the presentation of neoantigens or evading extrinsic induction of apoptosis. The detailed information on predicted neoantigens, cytolytic activity and FASLG of each sample is included in TableS4.

### Co-mutation of the proteasome and extrinsic apoptosis pathways and immune evasion in other cancer types

Co-mutation of the proteasome and extrinsic apoptosis pathways in the other five cancer types of the pan-cancer database is not enriched. Nevertheless, we investigated whether the trends we observed in colorectal cancer generalized to these five tumor types such that tumors with co-mutations in this pair of pathways were more likely to evade immune surveillance. The 1060 samples of these five cancers include 105 (9.9%), 97 (9.1%) and 196 (18.5%)samples with mutations in both pathways, the apoptosis pathway only and the proteasome pathway only, respectively. By one-way ANOVA, cases with co-mutation show significantly more predicted neoantigens (p-value < 2*10^−16^), higher levels of cytolytic activity (p-value = 0.02) and FASLG expression (p-value = 0.001) compared to the WT tumors across the five types of cancer.

### Clonality of pathway co-mutation

To determine whether mutations in both pathways co-occur in the same clone, we extracted the variant allele frequency (VAF) of mutations in 29 samples with mutations in both pathways from the colorectal cancer. There is no clear difference in the VAF in co-mutated genes within the proteasome (median=0.30) and apoptosis pathways (median=0.28, Figure 5E). Some of these tumors had multiple mutations within a pathway, and we represented the VAF of the pathway using the variant with maximal VAF. For the majority of tumors with co-mutated pathways, the maximum VAF of both pathways was similar within the tumor, indicating that these co-mutations occur within the same clone (Figure 5F). To exemplify this point, we applied clonal decomposition to one of the TCGA-A6-2676 samples using the Sciclone package(20) and identified four clusters of mutations. Mutations assigned to the same cluster co-occur in the same clone. We therefore infer that during progression of the tumor, the *PSMA4* pE181 del mutations in the proteasome pathway and *BID* p.D141N in the apoptosis pathway emerge in the same subclone that stems from a clone with a mutation in *TP53*. This suggests that mutations in the proteasome and apoptosis pathways are not driver events in colorectal cancer, but are acquired during cancer progression, allowing cancer cells to evade immune surveillance.

## Discussion

We developed and evaluated a patient-specific test for finding enriched co-mutated pathways in both simulated datasets and patient datasets from specific cancers and across several cancer types. Through permutation testing, we verified that our approach generated more accurate p-values than Fisher’s exact test (FET), a standard approach to identify significant co-occurrence events. Specifically, we demonstrated that our method achieved high precision in simulated data with either small or large variation in mutation load. We applied our method to mutation data of several cancers and found that co-mutated pathways such as the PI3K and IP3 pathways are prevalent across cancers. We also found that co-mutations in the proteasome and apoptosis pathways were enriched in colorectal cancer, but not in other tumor types. Furthermore, we showed that co-mutation of the proteasome and apoptosis pathways is associated with elevated levels of predicted neoantigens, cytolytic activity and decreased apoptosis.

When using FET, a large number of pairs of pathways were co-mutated. In FET, the assumption is that two events (mutations in two pathways) are independent of each other. However, this assumption is very likely to be violated if samples: a) have hyper-mutated phenotypes, b) are unevenly sequenced, and c) have variable tumor versus stromal cellularity. Specifically, genes are more likely to be mutated under a high overall tumor mutation load and mutations are less likely to be detected with low sequencing depth or large contamination of samples with normal or stromal cells. We therefore developed a patient-specific method to account for this variability. We showed that empirical p-values obtained by permutation tests, which required prolonged computational time, were consistent with p-values generated using our Poisson binomial model based approach, but correlated poorly with p-values obtained by FET.

By using spiked-in signals we simulated two datasets corresponding to scenarios with small or large variability in mutational load. As expected, both tests perform well in datasets with small variability in mutational load while FET over-estimated the significance of co-mutated pathways in datasets with large variability in mutational load between samples. In addition, we showed that the power to detect co-mutated events associated with a pair of pathways (*A* and *B*) in our patient-specific method depends on the mutation probability of pathway *A* and the conditional mutation probability of pathway *B* given the mutation status of pathway *A*. In all simulations, we used data that had been generated to have statistics similar to data from the Yale melanoma data set, demonstrating the value of our method in human cancer data sets.

Given the success of the method with simulated data, we subsequently applied our method to six datasets of different cancer types from TCGA as well as to the pan-cancer dataset, which is an aggregate of these six cancer datasets. We used the p-values without correcting for multiple hypothesis testing. We note however, that due to significant overlaps between pathways collected from different manually curated databases, application of false discovery rate (FDR) for correcting the p-values is invalid.

In the pan-cancer analysis, the most salient pair of co-mutated pathways was the PI3K and IP3 synthesis pathways. Co-mutation of these two pathways was observed across all cancer types. Previous studies have shown associations between PIK3CA mutations and PTEN mutations or between PIK3R1 mutations and PTEN mutations(7,27). By aggregating mutations at the pathway level, over 20% of the cancers had co-mutations in these two pathways. Co-mutation in the PI3K and IP3 pathways results in increased PIP3, which activates the AKT/mTOR pathway to sustain proliferative signaling, the most fundamental trait of cancer cells(28,29).

In colorectal cancer, we identified significantly co-occurring mutations in the proteasome and apoptosis pathways. Several reports have implied that mutations in the proteasome pathway or the apoptosis pathway play a role in evading immune surveillance (14,15), but to the best of our knowledge this is the first study which implies that co-mutation of these pathways, at least in colorectal cancer, may lead to enhanced immune evasion. We hypothesized that these two pathways synergistically protect tumors from the immune system as they are key for degradation and immune system presentation of neoantigens, and for immunity mediated apoptosis. To support this premise, we assessed the association between the mutation status and additional features of tumor cell survival. Mutations in proteins associated with the proteasome had a negative effect on cytolytic activity and tumors with alterations in the apoptosis pathway accumulated Fas ligand. More interestingly, after applying clonal deconvolution, we discovered that mutations in the proteasome and apoptosis pathways belonged to the same subclone during clonal evolution. Since the tumors were probed only once in one anatomic region, we could not infer which of these two pathways was altered earlier in the process of cancer progression. However, there are many more cases mutated only in proteasome pathway than in apoptosis pathway, indicating that mutations in the proteasome pathway may occur first. To test this hypothesis, it would require sequencing measurements at multiple time points from the same tumor. Sampling at multiple time points could also inform us whether a subclone with co-mutation proliferates faster compared than subclones without.

Cytolytic activity in samples where the proteasome and apoptosis pathways were co-mutated was higher than in the other three groups. A possible explanation is that a substantial subset of clones in tumors with co-mutation in the proteasome and the apoptosis pathways carry no mutations in the proteasome pathway. Due to the larger number of predicted neoantigens in samples with proteasome and apoptosis pathway co-mutations, this subset of clones is likely to be associated with a larger number of neoantigens compared with the number of neoantigens in the other three groups. Therefore, the subset of clones that carry no proteasome mutations in the tumors that have co-mutation in proteasome and apoptosis may present many of the neoantigens properly and display overall increased cytolytic activity. However, this can only be assessed by single cell experiments or clonal deconvolution of the predicted neoantigens.

In summary, we developed a new method to identify co-mutated pathways from population-wise cancer genomic data at the pathway level. Attempts to discover SNP interaction epistasis effects in GWAS have not been successful and currently there are no known observations of loci-loci or gene-gene interactions at the population level, even though the interactions exist at an individual level(30). This is mainly caused by the heterogeneous interactions and limited sample sizes. Similarly, there is high degree of heterogeneity in cancer and we tackled this by integrating mutations at the pathway level. While we focused initially on mutational data, this method can be extended to other types of genomic abnormalities such as CNV, DNA methylation, etc. Due to the heterogeneity between tumors, we expect to capture more low frequency co-mutated pairs of pathways by integrating alterations of different levels. Another future direction is to validate our findings using single cell RNA sequencing, inspecting whether co-mutated transcripts are expressed within single cell. This direct measurement does not rely on deconvolution methods such as Sciclone which can often have non-unique solutions. Furthermore, inferences of order in which mutations occur first during cancer evolution are superior to inferences based on deconvolution approaches. Detection of co-dysregulated processes and the order in which they evolve can inform treatment decisions in individual patients and advance personalized medicine.

## Acknowledgement

This research was partially funded by NIH grant 1R01HG008383-01A1 (Y.K.).

---

**Figure S1:**
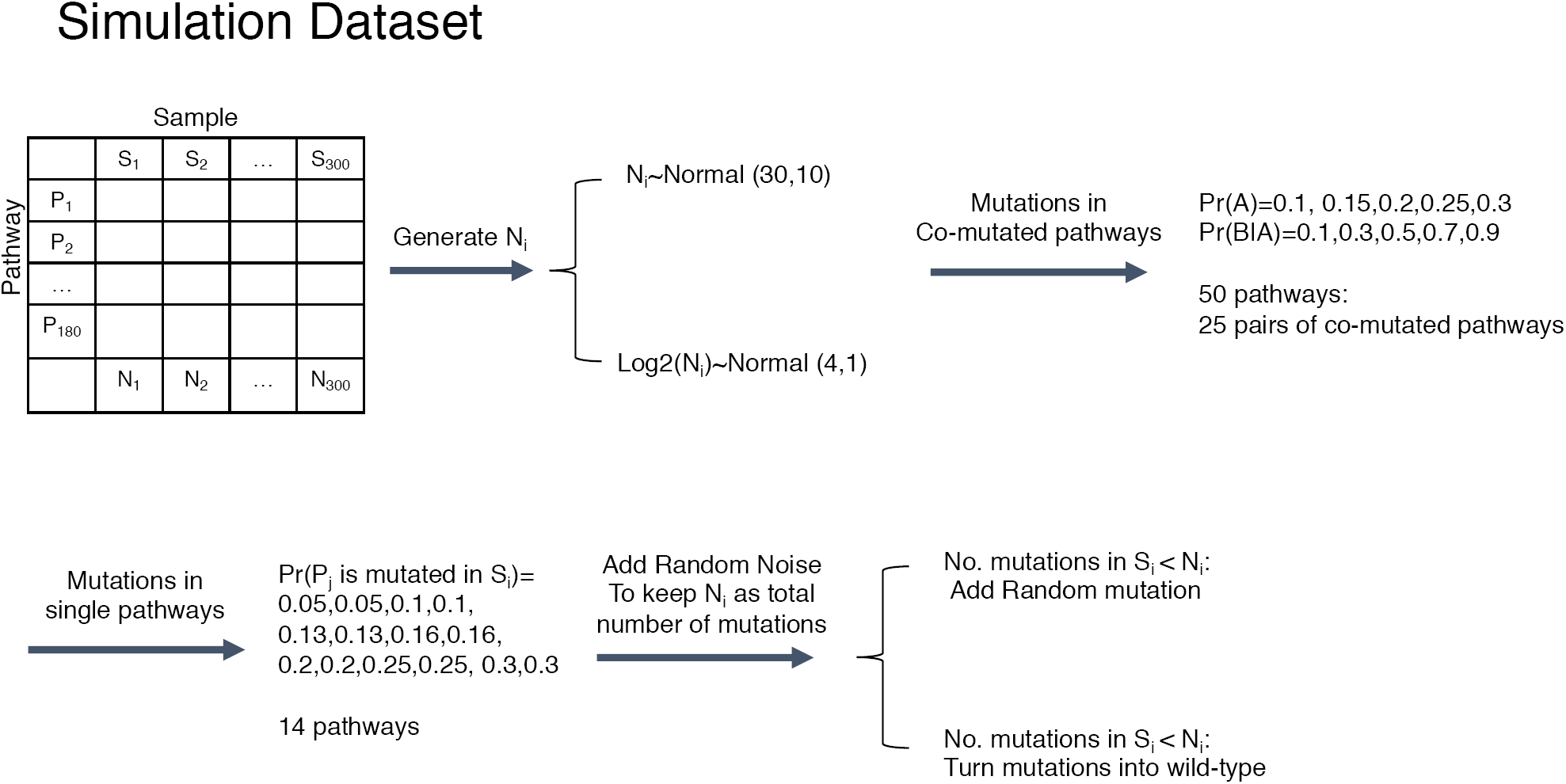
Procedure for generating simulated datasets

**Figure S2:**
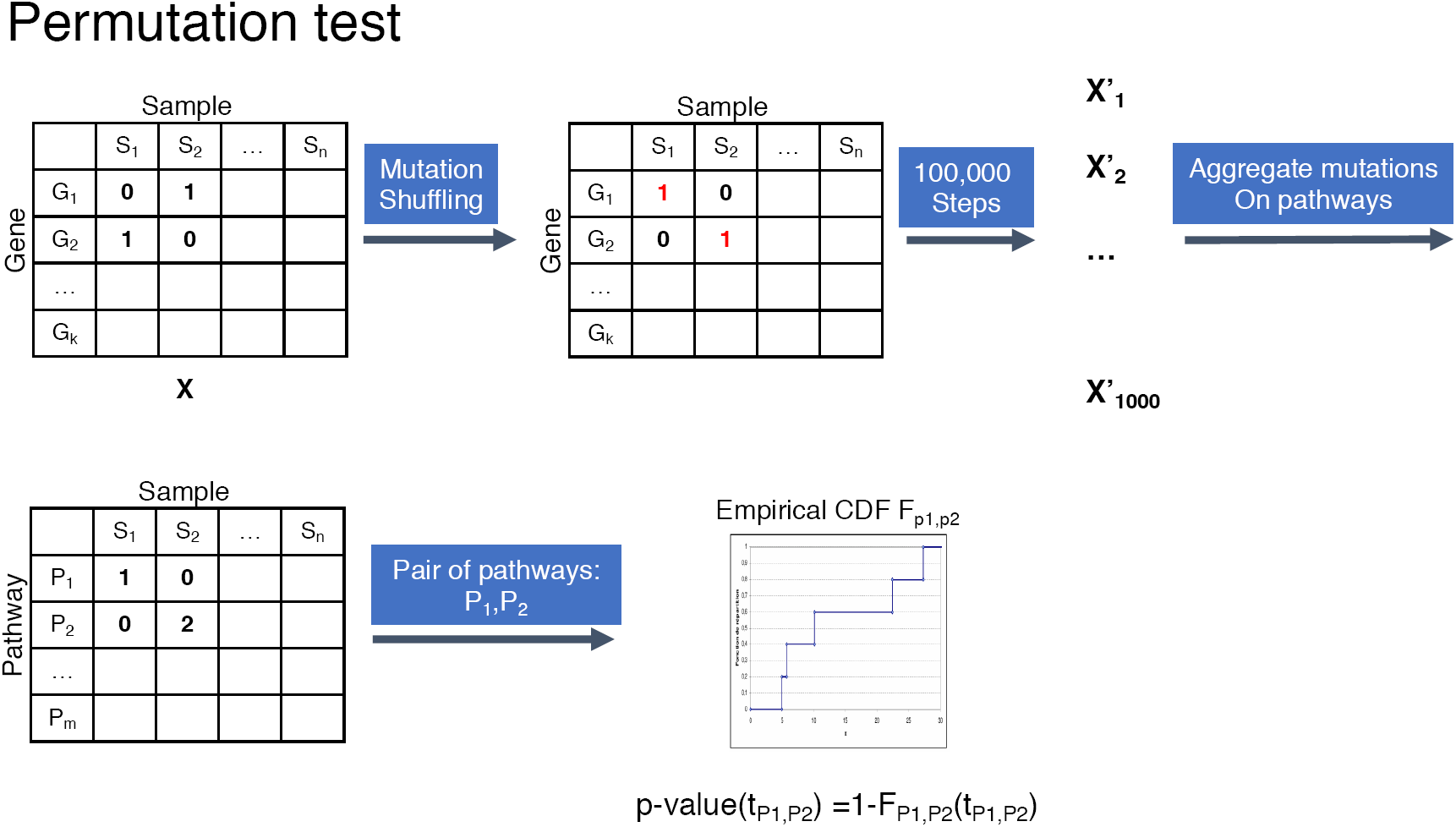
Procedure for generating pathway co-mutation p-values based on permutation

**Figure S3:**
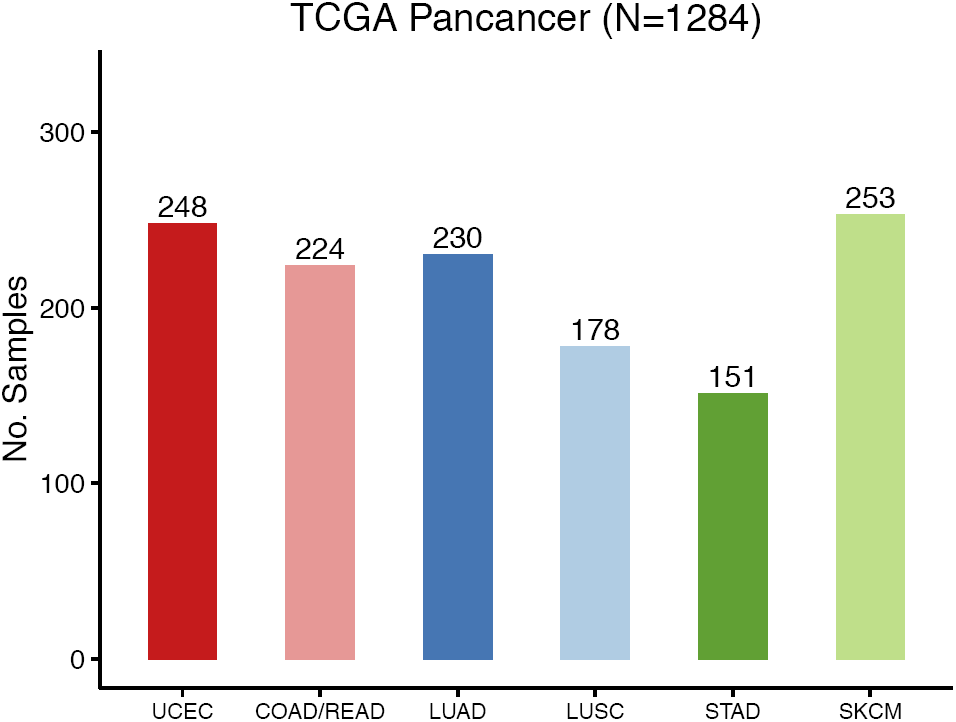
Sample size of six different type of cancer from the TCGA project

**Figure S4:**
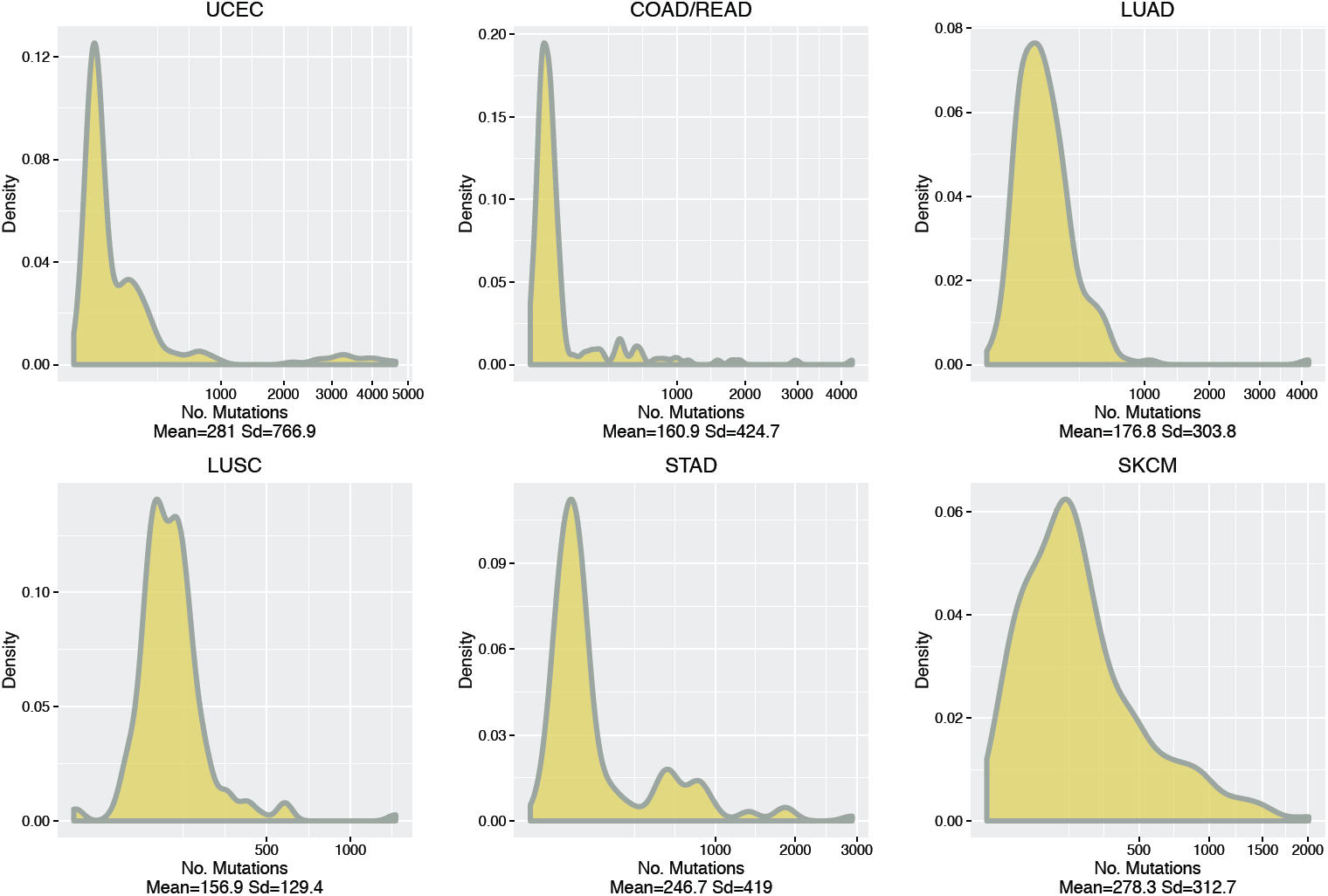
Distribution of number of mutations in six different types of cancers from the TCGA project

